# Phenotypic extremes or extreme phenotypes? On the use of large and small-bodied “phenocopied” *Drosophila melanogaster* males in studies of sexual selection and conflict

**DOI:** 10.1101/2022.07.27.501796

**Authors:** Kyle Schang, Renée Garant, Tristan A.F. Long

## Abstract

In the fruit fly, *Drosophila melanogaster*, variation in body size is influenced by a number of different factors and may be strongly associated with individual condition, performance and success in reproductive competitions. Consequently, intra-sexual variation in size in this model species has been frequently explored in order to better understand how sexual selection and sexual conflict may operate and shape evolutionary trajectories. However, measuring individual flies can often be logistically complicated and inefficient, which can result in limited sample sizes. Instead, many experiments use large and/or small body sizes that are created by manipulating the developmental conditions experienced during the larval stages, resulting in “phenocopied” flies whose phenotypes resemble what is seen at the extremes of a population’s size distribution. While this practice is fairly common, there has been remarkedly few direct tests to empirically compare the behaviour or performance of phenocopied flies to similarly-sized individuals that grew up under typical developmental conditions. Contrary to assumptions that phenocopied flies are reasonable approximations, we found that both large and small-bodied phenocopied males frequently differed from their standard development equivalents in their mating frequencies, their lifetime reproductive successes, and in their effects on the fecundity of the females they interacted with. Our results highlight the complicated contributions of environment and genotype to the expression of body size phenotypes and lead us to strongly urge caution in the interpretation of studies solely replying upon phenocopied individuals.

## Introduction

Within populations, there frequently exists - often considerable - variation between individuals in the size of their bodies. Putting aside the obvious source of variation associated with differences in the age of conspecifics (*see* Ellstrand 1983), understanding the causes and consequences of such intra-population variation is of great interest to biologists (Blanckenhorn 2000). This is because variation between individuals is an essential pre-requisite for natural and/or sexual selection to operate, as if everyone in a population expressed the same phenotypes, any individual differences in performance would be due to stochastic factors, rather than attributable to competitive advantages associated with their traits (Darwin 1859, Gregory 2009). In fact, variation in body size is often ubiquitous as it reflects variation at loci across the genome (*e.g*. Oldham *et al*. 2000, Turner *et al*. 2011) and/or the contributions of the developmental environment (*reviewed in* Mirth and Shingleton, 2012). Furthermore, variation in body size in populations is strongly correlated with individual variation in fitness in a wide range of taxa, with those of larger body sizes frequently at a greater fitness advantage compared to smaller individuals (Partridge and Farquhar 1983, Savalli and Fox 1998, Sokolovska *et al*. 2000, *but see* Blanckenhorn 2000). However, defining “one of the most important quantitative traits under evolutionary scrutiny” (Blanckenhorn 2000) can be logistically challenging, as body size is a composite trait (*aka* a “bloodless geometric construct” Bonner 2011), and studies of body size must involve subjective decisions about what specific variables will be used as indices.

In fruit flies (*Drosophila melanogaster*), a model species used in a wide range of genetic, behavioural, sexual selection and sexual conflict studies (Weiner 2000, Brookes 2001), the importance of body size variation is frequently explored, and is quantified in a number of different ways. Flies may be weighed individually on a microbalance scale (*e.g*. Azevedo 1997, Klok *et al*. 2009), or size might be inferred by measuring wing length (*e.g*. Reeve *et al*. 2000), thorax length (*e.g*. Ewing 1961, Partridge and Farquhar 1983, Partridge *et al*. 1987) and/or total body length (e.g. Morgan *et al*. 2016) of flies viewed through a microscope. Measuring body size using these techniques can be logistically difficult, and requires considerable investments of time, effort and patience. This may ultimately be a limiting factor in obtaining a sufficient number of flies of a desired body-size phenotype for experimental studies, possibly limiting the study’s sample size and the statistical power of the analyses. As such, it has become fairly common practice to artificially obtain large or small flies via the manipulation of larval diet and/or the degree of larval competition via a process called “phenocopying” (*sensu* Goldschmidt 1945) in which the development of individuals resembling a desired phenotype can be induced via environmental manipulation. It has long been known that reducing the available adult resource pool (Mueller *et al*. 1993) by increasing larval densities to heighten competition for resources (or alternately by decreasing the amount of nutrition in the developmental environment) will result in eclosed adult flies of relatively small body size (Beadle *et al*. 1938, Miller and Thomas 1958, Ewing 1961, Miller 1964), while larger than average adults can be produced by ensuring ample resources during their larval phase. An alternative (and less frequently used) method or producing flies of different sizes involves altering developmental temperature (*e.g*. Zamudo *et al*. 1995). In *D. melanogaster*, like other holometabolous insects, adult body size is largely determined by the amount of resources that are acquired during the larval phase (Boggs 1981). The technique of phenocopying via nutritional manipulation thus offers a quick and convenient method of obtaining flies of large- or small-bodied phenotypes, allowing biologists to compare the individuals divergent in “condition” (defined as the pool of resources an individual has available to invest in their trait expression *sensu* Rowe and Houle 1996).

Implicit in their use is the assumption that the resulting groups of phenocopied flies are for all extents and purposes equivalent to the large- or small-bodied flies that one would be able to (eventually) collect using the more onerous methods described above. However, there are several reasons to be concerned about the validity of that assumption. First, experimentally manipulating larval densities may change the intensity and/or nature of selection acting on them, as increased larval densities leads to more intra-specific competition, to decreased and degraded nutritional resources, and increased exposure to environmental waste products (Borash *et al*. 1998), which can result in lower survivorship (Khodaei *et al*. 2020), slowed developmental rates, and worsened individual condition (*reviewed in* Ashburner *et al*. 2005, Borash *et al*. 1998, Khodaei and Long 2019). Furthermore, the developmental conditions created by changing the available nutrition (either directly, or indirectly via changes in larval density) may result in changes to the selective environment that are dramatically different than those that flies have typically experienced (and presumably had the opportunity to adapt to). In populations maintained at high larval densities, selection alternately favours different genotypes associated with larval feeding rates and tolerance to waste products depending on the extent of nutritional depletion and ammonia accumulation in their environment (Borash *et al*. 1998). Similarly, Burns *et al*. (2012) observed that the effects of nutritional restriction on the expression of exploratory behaviour and fecundity depended on a fly’s genotype, suggesting that phenotypic plasticity arising from genotype-by-environment interactions (GxE) in novel developmental environments is another unintentional consequence associated with the use of phenocopying technique. Together, it is likely that the genetic diversity and composition of the surviving adult flies that are obtained via phenocopying will be different than those that emerge under normal developmental conditions. Thus, if an individual’s body size is the product of both their genetic characteristics (Oldham *et al*. 2000; Turner *et al*. 2011), and their developmental history (Beadle 1938, Miller and Thomas 1958, Boggs 1981) it may not be logical to assume that two flies sharing similar body size, yet arriving at that “same” phenotype via different pathways, would be equivalent in their physical capabilities and behaviours.

Surprisingly, a review of the literature failed to reveal any systematic empirical tests of this key assumption (*with the notable exceptions of* Ewing 1961, and Verma *et al*. 2022). In this study, we set out to experimentally examine whether these phenocopied flies are representative of those reared under normal conditions by comparing male flies of the same body size, yet of different developmental histories. We focused our attention on male reproductive performance and its consequences as fruit flies are often used in studies examining aspects of sexual selection and sexual conflict. We did so by performing assays measuring behaviours, reproductive successes and the effects of these males on their mates to determine if any differences exist between phenocopied flies and those reared under normal culturing conditions that they are meant to represent.

## Materials & Methods

### Population Origins & Typical Culturing Protocols

All flies used in our experiments were derived from the “*Ives*” (hereafter “IV”) population of *Drosophila melanogaster*. This population was founded from a sample of 200 male and 200 female flies that were collected in the vicinity of South Amherst, MA (USA) in 1975, and have been cultured following a standardized protocol since 1980 (Rose 1984, Tennant *et al*. 2014). This is a large (~3500 adults/generation), outbred wild-type population that is maintained in 25×95 mm vials, on non-overlapping 14-day generations, at 25°C, at 60% relative humidity and on a 12L:12D diurnal cycle. At the start of each generation, adult flies are removed from all vials, mixed *en masse* under light CO_2_ anesthesia and distributed amongst 35 vials, that each contain ~ 10mL of banana/agar/killed-yeast media as a light sprinkle of live yeast. Flies oviposit in these vials for up to 24h before being removed and the number of eggs in each vial is trimmed by hand to a standard density (100 eggs/vial) before being returned to the incubator.

In addition to flies from the IV population, we also used flies from the “sister” IV-*bw* population that was created by introgressing (via repeated rounds of back-crossing) the recessive *bw^1^* allele into the IV genetic background. Flies in this population exhibit the homozygous brown-eyed phenotype (instead of the dominant red-eyed *wild type* phenotype) necessary to determine paternity of offspring for experiment 1, but are otherwise identical to the IV population, and are maintained following the same protocols.

### Phenocopying and size sorting protocols

The “target” IV flies used in our experiments were obtained from eggs that had been laid by placing mated adult females into half-pint containers fitted with small petri dishes containing an agar/grape-juice media and a smudge of live yeast at the opening of each container. These petri dishes were removed ~ 16h later and sets of precisely counted eggs were carefully removed from the surface of the media and transferred to vials containing 10mL of the banana-based media. In our experiments we created our experimentally small/large flies by altering larval density by transferring over greater/fewer numbers of eggs into the vials than the 100/eggs/vial that is typical of our populations’ culture protocol. To produce the experimentally large flies, vials were seeded with 50 eggs each, while to produce experimentally small flies, vials received sets of 200 eggs apiece. At the same time we also created vials containing 100/eggs each using eggs collected from the IV population. These vials were used to create control large/small males (using the protocol described below), for use in both our sets of experiments, as well as the female flies used in our second experiment. When conducting our first experiment, we also created vials containing 100 similarly-aged eggs each obtained from the IV-*bw* population. All vials were incubated under standard environmental conditions, and starting 8-9 days later, we collected adult virgin flies (within 8h of their eclosion from pupae), which were separated by sex and stored in fresh vials in groups of ~40 for 1-3 days prior to the start of our assays.

All of our assays involved comparisons of the performance and/or success of large- and small-bodied flies. To obtain flies in these categories we used the high throughput sieve-sorting protocol described in Long *et al*. (2009). Briefly, lightly anesthetized flies are placed at the top of a column of electro-formed sieves (Precision Eforming, Cortland, NY, USA) mounted on a Gilson Performer SS-3 3” sieve shaker (Gilson Company Inc., Lewis Center, OH, USA). The sieves range in hole diameter from 887μm to 1420μm, with the diameter of the holes decreasing by ~5% at each layer down the column of sieves. The anesthetized flies are agitated at a rate of 3,600 vibrations min^−1^ for a total of 4 min. For all experiments, “small” male flies were defined as those that were small enough to pass through the 1167μm diameter sieve, whereas “large” flies were those that were too large to pass through the 1365μm diameter sieve. Both the large and small control male flies were collected in this manner from the vials initiated with sets of 100 eggs, while the experimental large male flies were those males ≥ 1365μm from the vials initiated with sets of 50 eggs, and the experimental small male flies were those ≤1167μm from the vials initiated with sets of 200 eggs. Once sorted, flies were kept in fresh vials overnight to allow them time to recover from their anesthesia/sieving experiences prior to the start of the assay.

### Experiment 1: Mating success assay

In our first assay we set out to compare the behaviour and subsequent reproductive success of single similarly-sized phenocopied or control males in a competitive environment. The assay began by placing a single “focal” IV male (belonging to one of the 4 different types, with 49-50 replicate vials/type) into vials containing 10mL of media and 3 “competitor” IV-*bw* males and 3 IV-*bw* females starting ~9:00am on Day 10 of the flies’ culture cycle. Vials were immediately placed horizontally on an observation board in a well-lit room and monitored constantly for the next 120 minutes. All matings between the focal male and any of the IV-*bw* females was recorded. After the 2h observation period had elapsed, flies were immediately anesthetized and separated by sex. Females were retained in the observation vial, and males were placed into a fresh vial. The following day at 9:00am the females were transferred into the male’s vials for an additional 2h observation period before being separated overnight. This procedure was repeated for a total of five consecutive days, with flies discarded at the start of the 6^th^ day. All observation vials were retained, and incubated for 14 days at which time the number of adult brown-eyed flies (sired by the competitor males) and red-eyed flies (sired by the wild-type focal male) were counted.

### Experiment 2: Male exposure assay

In our second assay, we set out to measure the effects of *D. melanogaster’*s antagonistic male persistence (e.g., the unrelenting courtship and repeated mating attempts of females, *reviewed in* Long *et al*. 2009) for similarly-sized phenocopied and control males. We did so by comparing the effects of short vs long-term exposure to these males of on the fecundity of groups of females, a well-established method of quantifying this outcome of sexual conflict (*see* Rice et al. 2006, Filice and Long 2016). We began by creating 240 vials each containing 10 adult virgin IV females, which were haphazardly divided into 4 groups. Starting around 9:00am on Day 10 of the flies’ culture cycle we added (under light anesthesia) 10 males belonging to one of the 4 different types into the vial. These vials were placed horizontally on an observation board in a well-lit room for a period of 3 hours. At that time, half the vials from each group were haphazardly selected for the “short-exposure treatment”. Males from these vials were removed and the females in the vial were returned to the observation board. The vials that were not selected thus belonged to the “long-exposure treatment”. For the next 3 days these long-exposure treatment vials were regularly scanned for observations of matings (12 observation sessions on the 1^st^ day, 14 sessions on the 2^nd^ day, each spaced approximately 30 minutes apart plus a single observation session on the 3^rd^ day). Since there were no males present in the short-exposure treatment vials, no behavioural observations were performed. On the 3^rd^ of the assay, all females from all 4 treatments (30 replicate vials/treatment) were removed from their vials, and placed individually into 13×100 mm test tubes containing 2mL of the banana-based media (with a scored surface to encourage oviposition) for a period of 24h before being discarded. These test-tubes were incubated for 14 days, and the number of adult offspring present were counted. We then calculated the mean offspring production across the 10 test-tubes yielded by each vial.

### Statistical Analyses

All data analyses were conducted in the R statistical computing environment (version 4.0.3, R Core Team, 2020). We compared the cumulative number of matings observed in our first experiment by large or small control males and their phenocopied male counterparts using generalized linear models (GLMs), with quasipoisson error distributions. Similarly, we analyzed the number of wild-type offspring (sired by the focal males) eclosing from the first days’ vial, as well as cumulatively over the 5 days of the experiment with GLMs with quasipoisson error distributions. The significance of the independent factors in our GLMs was determined using the *Anova* function from the *car* package (Fox and Weisberg 2011). The magnitude of the difference between the control flies and the phenocopied experimental flies was quantified using the Cliff’s delta effect size statistic using the *cliff.delta* function in the *effsize* package (Torchiano 2020).

For our second assay’s data we compared the median number of offspring produced by females in vials that differed in the type of male they were exposed to (control or phenocopied), the length of their exposure to males (3h or 3days), and their interaction. As the assumption that each sub-group was normally distributed and/or had homogenous variances was violated, instead of conducting a 2-way ANOVAs, data were analyzed using the non-parametric Scheirer-Ray-Hare method (Sokal and Rohlf 1995), implemented with the *scheirerRayHare* function in the *rcompanion* package (Mangiafico 2021). In the case of statistically significant interactions, group medians were compared with Dunn’s test (Dunn 1964) with Holm’s (1979) multiple-comparison correction method using the *dunn.test* function in the eponymous R package (Dinno 2017).

Using the second experiment’s behavioural data collected from the flies in the long-exposure treatment vials, we compared the total number of copulations observed across all 3 days/27 sessions for the large and small males against their experimental phenocopied counterparts using a GLM with quasipoisson errors. The magnitude of the difference between these groups was also estimated using Cliff’s delta method.

## Results

### Experiment 1: Mating success assay

Over the course of five days’ observation periods we observed no statistically significant difference in means of the total number of copulations involving focal male flies that had developed under standard larval densities and size-matched phenocopies (Large male type treatment, GLM: LLR *χ*^2^= 1.49, df= 1, p=0.223, Cliff’s delta (±95%CI): −0.146 (−0.315, 0.031), *Figure 1a*; Small male type treatment, GLM: LLR *χ*^2^= 0.30, df= 1, p= 0.587, Cliff’s delta (±95%CI): 0.037 (-0.155, 0.226), *Figure 1b*).

**Figure 1:**
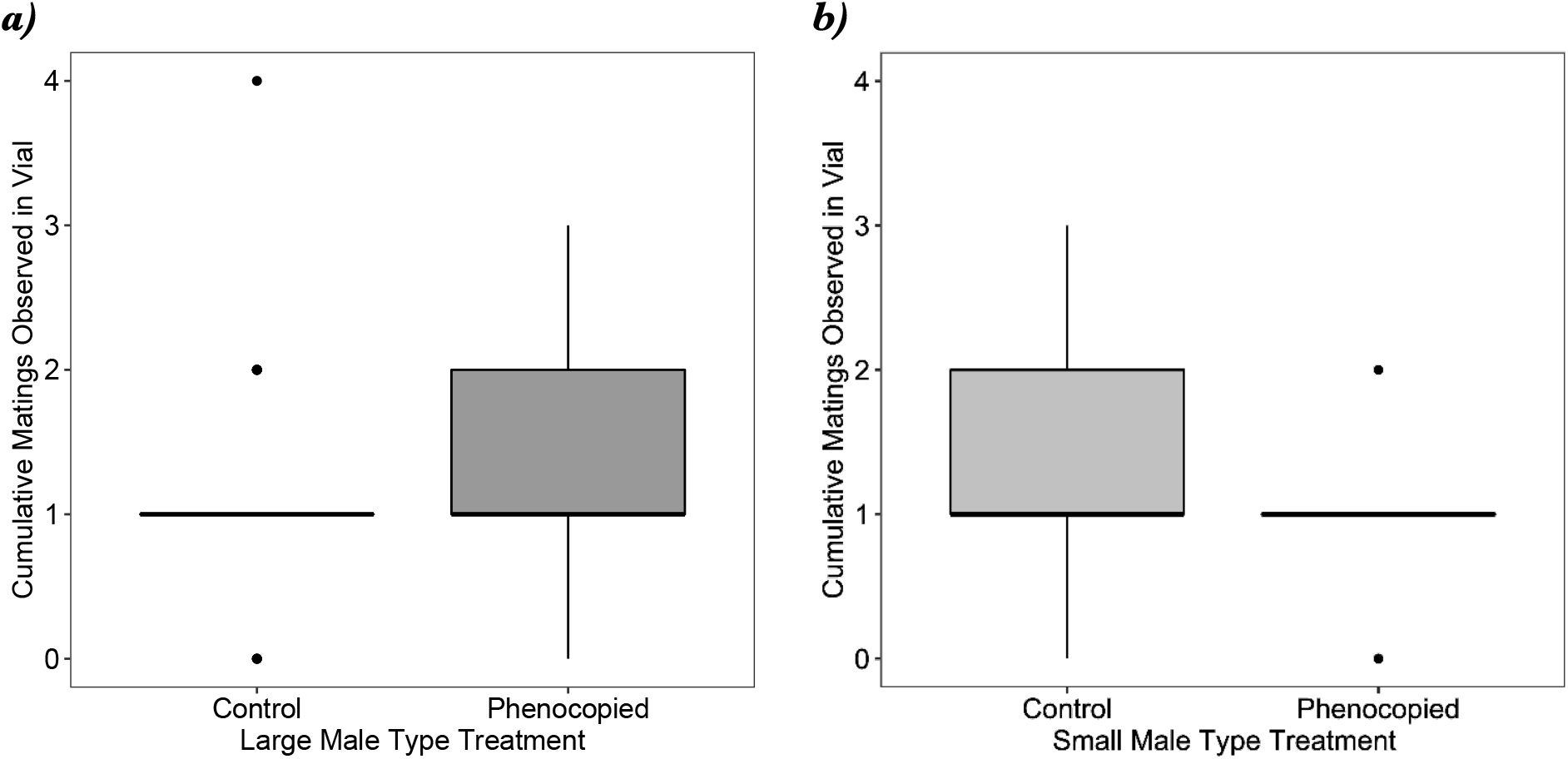
Boxplots showing the total number of copulations observed involving single focal male fruit flies, *Drosophila melanogaster*, housed in a reproductively competitive environment observed over five consecutive days. Focal flies were either large (Figure 1a) or small (Figure 1b), and achieved that size either under standard developmental conditions (control), or as a result of environmental manipulation (phenocopied). The boxes enclose the middle 50% of data (the inter-quartile range, IQR), with the thick horizontal line representing the location of median. Data points >± 1.5*IQR are designated as outliers. Whiskers extend to largest/smallest values that are not outliers, indicated as closed circles.

In total we counted 11,505 red-eyed offspring (sired by the focal males). When comparing the number of offspring sired by the focal males, we saw in the large male type treatment, that phenocopied males were modestly (but not significantly) more successful than control males (GLM: LLR *χ*^2^=1.66, df= 1, p= 0.198, Cliff’s delta (±95%CI): −0.169 (−0.379, 0.057) *Figure 2a*), a trend that increased slightly when examining the cumulative offspring production over five days (GLM: LLR *χ*^2^= 3.09, df= 1, p= 0.08, Cliff’s delta (±95%CI): −0.233 (−0.440, −0.002), *Figure 2c*). For the focal males in the small male type treatment, we saw that phenocopied males sired more offspring than control males both on the first day of the assay (GLM: LLR *χ*^2^= 6.49, df= 1, p= 0.010, Cliff’s delta (±95%CI): −0.362 (−0.553, −0.136), *Figure 2b*) as well as cumulatively over the five days of the assay (GLM: LLR *χ*^2^= 9.414, df= 1, p= 0.002, Cliff’s delta (±95%CI): −0.349 (−0.539, −0.125), *Figure 2d*).

**Figure 2:**
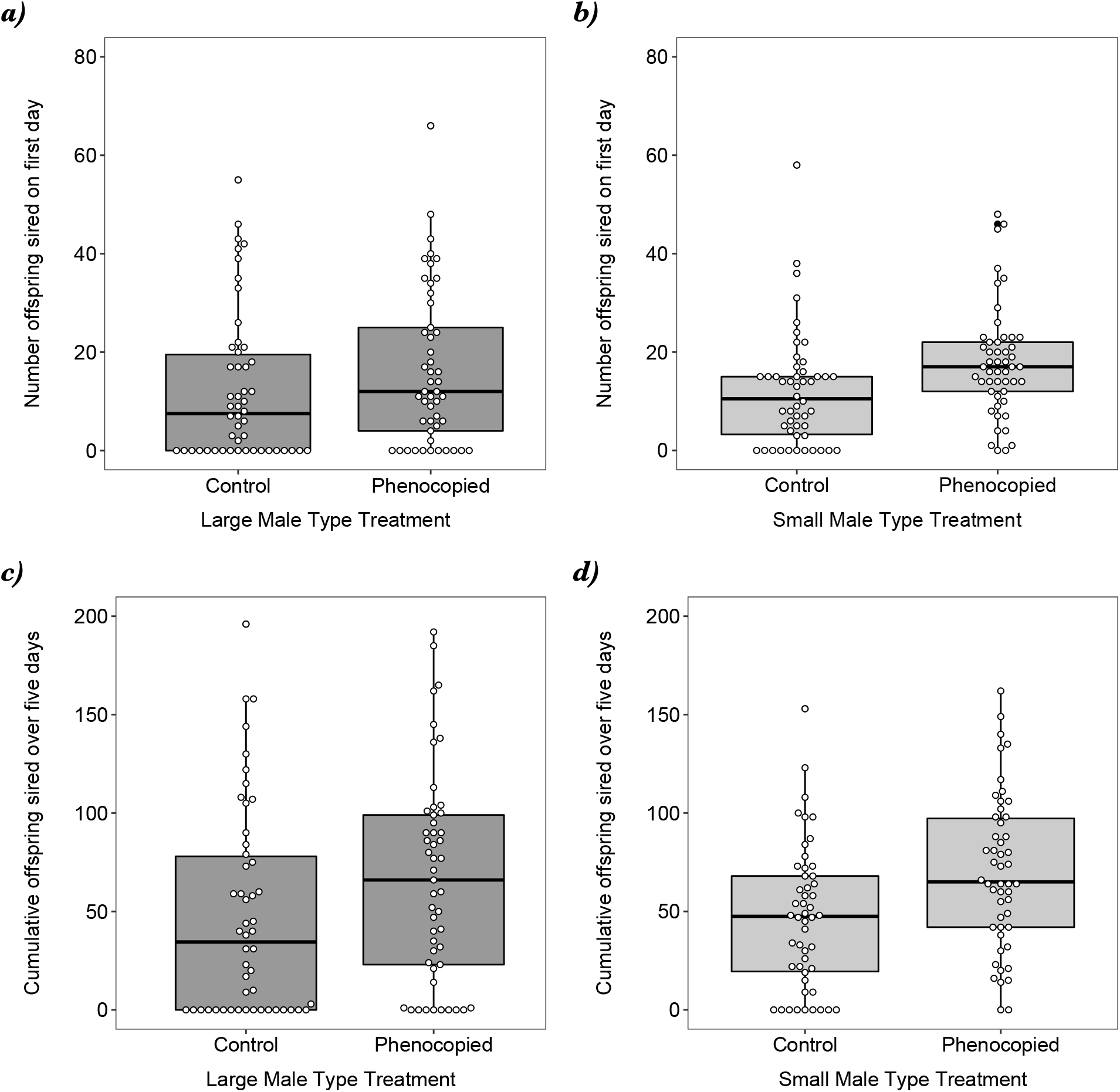
Boxplots showing the number of offspring sired by single focal male fruit flies, *Drosophila melanogaster*, housed in a reproductively competitive environment after one day (Figures 2a & 2b), or cumulatively over five consecutive days (Figures 2c & 2d). Focal flies were either large (Figures 2a & 2c) or small (Figures 2b & 2d), and achieved that size either under standard developmental conditions (control), or as a result of environmental manipulation (phenocopied). The boxes enclose the middle 50% of data (the inter-quartile range, IQR), with the thick horizontal line representing the location of median. Data points >± 1.5*IQR are designated as outliers. Whiskers extend to largest/smallest values that are not outliers. Values of datapoints indicated with open circles.

### Experiment 2: Male exposure assay

For the vials in the long exposure treatment, we counted the number of copulations in 27 sessions spanning 3 days. For those females housed with large-bodied males (*Figure 3a*) we saw no difference in the mean number of cumulative matings in the control and the phenocopied groups (GLM: LLR *χ*^2^= 0.24305, df= 1, p= 0.622, Cliff’s delta (±95%CI): 0.076 (−0.216, 0.355)). In contrast, in the vials where males were small (*Figure 3b*), there were markedly fewer matings observed involving phenocopied males compared to control males (GLM: LLR *χ*^2^= 11.417, df= 1, p= 7.28 × 10^−4^, Cliff’s delta (±95%CI): 0.49 (0.206, 0.698)).

**Figure 3:**
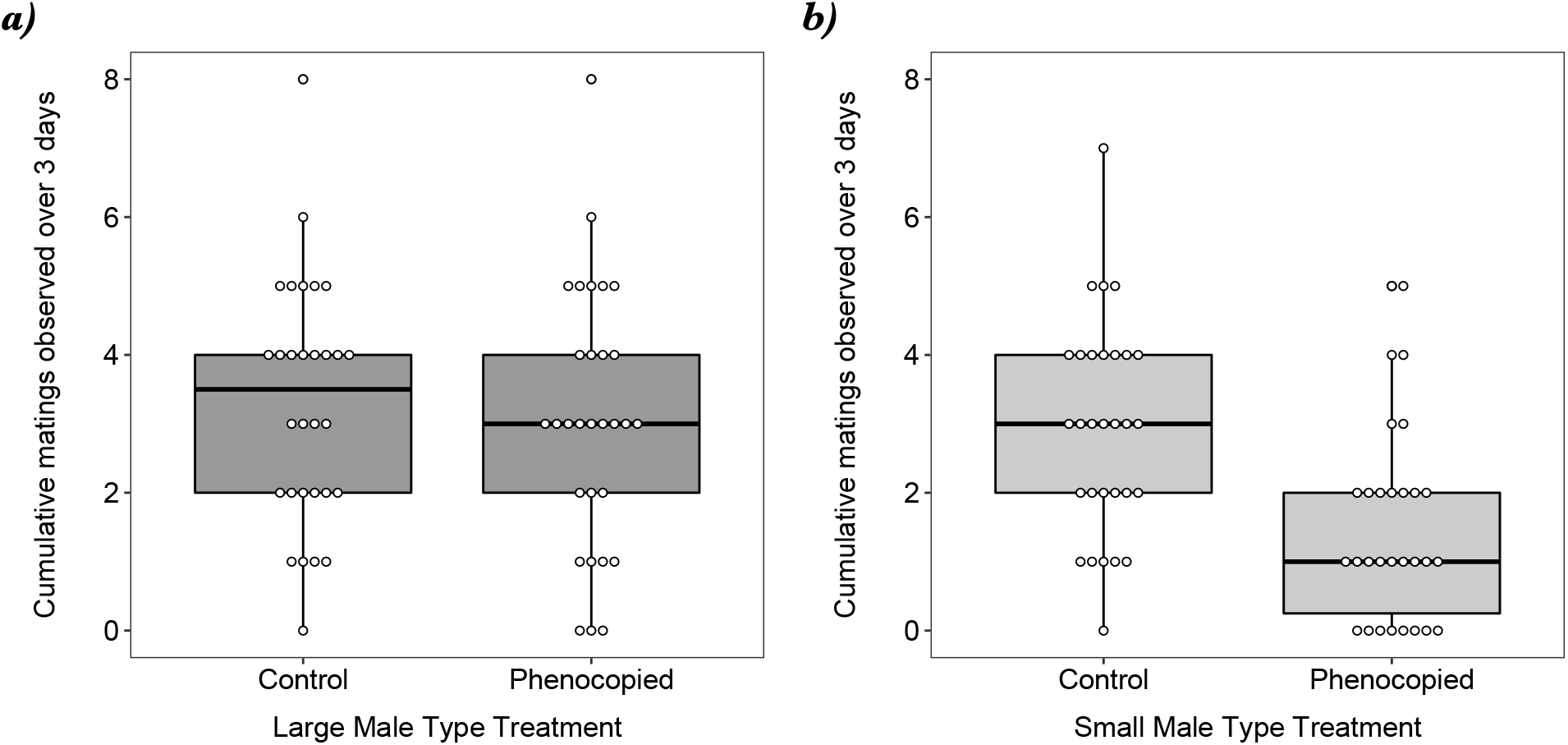
Boxplots showing the total number of copulations observed involving a group of male fruit flies, *Drosophila melanogaster*, housed in a reproductively competitive environment over 3 days’ worth of observation sessions. Focal flies were either large (Figure 3a) or small (Figure 3b), and achieved that size either under standard developmental conditions (control), or as a result of environmental manipulation (phenocopied). The boxes enclose the middle 50% of data (the inter-quartile range, IQR), with the thick horizontal line representing the location of median. Data points >± 1.5*IQR are designated as outliers. Whiskers extend to largest/smallest values that are not outliers, indicated as closed circles.

In total, we counted 52, 213 offspring produced by females in this assay. For females that were exposed to large bodied males (*Figure 4a*), we observed significant differences in mean offspring production associated both with the type of male, and the duration of exposure, but no significant interaction between these factors (*Table 1a*). Females exposed to groups of control large males produced more offspring overall (median= 23.79) than those that were exposed to phenocopied males (median= 20.72) (Cliff’s delta (±95%CI): 0.426 (0.225, 0.592)). Females also produced more offspring if their exposure to males has been brief (median = 23.94) than they did with a prolonged exposure (median = 21.06) (Cliff’s delta (±95%CI): −0.404 (−0.568. −0.210)). For females that were exposed to small bodied males (*Figure 4b*), we saw that offspring production depended on both the type of male (control or phenocopied), as well as the duration of the exposure (*Table 1b*), with the magnitude of the difference between long- and short-exposure much more pronounced when the males were from the control group compare to when they originated from the phenocopied group.

**Figure 4:**
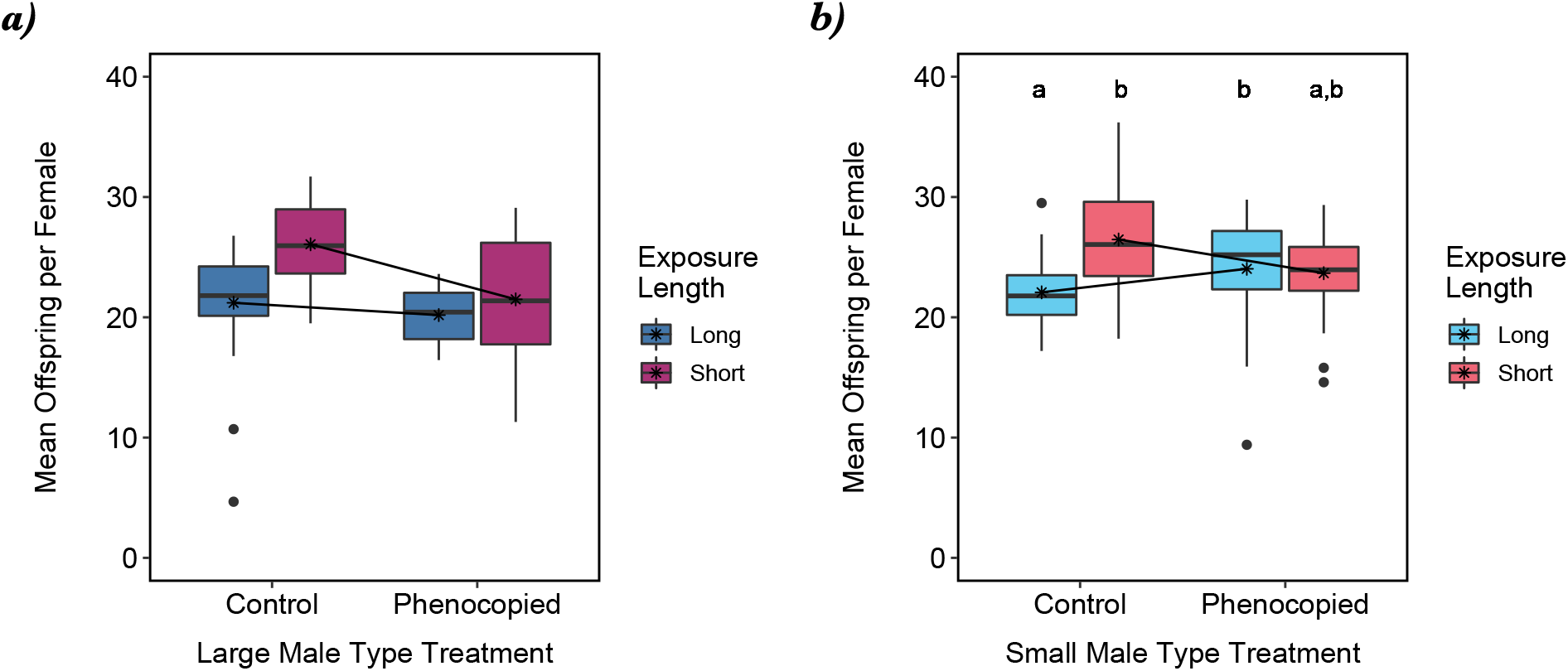
Boxplots showing the mean number of offspring produced by female fruit flies, *Drosophila melanogaster*, that were exposed to groups of males for a period of 3 hours (“Short”) or 3 days (“Long”). Groups of males flies were either large (Figure 4a) or small (Figure 4b), and achieved that size either under standard developmental conditions (control), or as a result of environmental manipulation (phenocopied). The boxes enclose the middle 50% of data (the inter-quartile range, IQR), with the thick horizontal line representing the location of median. Data points >± 1.5*IQR are designated as outliers. Whiskers extend to largest/smallest values that are not outliers, indicated as closed circles. The location of each group mean is indicated by a star. In Figure 4b the results of a post-hoc Dunn’s test comparing group medians is indicated by letters, where groups that do not share the same letter are considered statistically different.

**Table 1.** Results from Scheirer–Ray–Hare analyses on the mean offspring production of female fruit flies, *Drosophila melanogaster*, that were exposed to groups of either a) large bodied or b) small bodied male flies. Groups of male flies reached their body size either under standard developmental conditions, or as a result of environmental manipulation, and the exposure period was either 3 hours or 3 days.

| **Male Size** | **Factor** | **df** | **SS** | **H** | **p** |
| --- | --- | --- | --- | --- | --- |
| **a) Large** | Male Type | 1 | 19610 | 16.208 | 5.7 x 10<sup>-5</sup> |
|  | Exposure Length | 1 | 17642 | 14.581 | 1.3 x 10<sup>-4</sup> |
|  | Male Type x Exposure Length | 1 | 3532 | 2.919 | 0.088 |
|  | Residual | 116 | 103190 |  |  |
| **b) Small** | Male Type | 1 | 140 | 0.118 | 0.732 |
|  | Exposure Length | 1 | 6407 | 5.384 | 0.020 |
|  | Male Type x Exposure Length | 1 | 13505 | 11.349 | 7.5 x 10<sup>-4</sup> |
|  | Residual | 115 | 120358 |  |  |

## Discussion

Body size is a complex composite trait that is influenced by both genetic and environmental factors. Variation in body size is often associated with dramatic differences in performance (Partridge and Farquar 1983), as it can be linked to an individual’s ability to acquire resources, compete for mates and produce offspring. In fruit flies, *Drosophila melanogaster*, body size is frequently connected to many traits that are directly or indirectly associated with condition or fitness variation. The basis for these connections, however, are often based on observations using large- or small-bodied “phenocopied” flies, rather than flies that had arrived at those extreme phenotypes under typical developmental conditions. In this study we set out to test the assumption that large- and/or small-bodied “phenocopied” male fruit flies had similar behaviours, reproductive successes, and effects on their mates as those that they are meant to approximate. Our results suggest that this method to obtain flies at the phenotypic extremes of body-size distributions yields flies that are superficially similar, but are ultimately quite different than the flies they are supposed to represent.

In our assays we frequently observed meaningful differences between phenocopied and control flies in their behaviours, their reproductive successes and/or their effects on females, and that the nature of these differences varied with the body size category under examination. First, when considering large-bodied flies, we did not detect significant differences in the mean number of copulations involving phenocopied or control flies in either our first or second experiments (*Figures 1a & 3a*). Similarly, in the first experiment, the mean reproductive successes of individual male flies from the two groups were not dramatically or significantly different from each other (*Figures 2a & 2c*). In our second assay, where we manipulated both male origin and exposure treatment, we saw a negative effect of prolonged male exposure on female fecundity that is consistent with other studies (*e.g*. Lew and Rice 2005, Filice and Long 2016). However, we also detected a significant effect of the type of male on female fecundity, with females exposed to control flies producing ~ 14% more offspring than those exposed to phenocopied males (*Figure 4a*). When focusing on small-bodied flies, while we did not observe any significant differences between groups in the number of matings in our first assay, the phenocopied flies were more successful at siring offspring than were control males (*Figures 2b & 2d*). Interestingly, in our second assay, where groups of flies were housed with groups of females for a period of 3 days, the phenocopied males were observed mating less frequently than control males. When the offspring production of females in our second assay were analyzed we detected a significant interaction between male type and exposure length: for control flies, longer exposure was associated with lower productivity, but for phenocopied flies, there was no significant difference between groups. Taken together these results lead us to reject the assumption that the condition and/or reproductive abilities of phenocopied male flies are equivalent to individuals found at the phenotypic extremes of a population’s body size distribution. While our study only used one possible method of phenotyping (manipulation of larval density), we suspect that flies of large- and/or small-body size produced by altering developmental nutrition or temperature may also prove to be imperfect mirror-images.

To the best of our knowledge, there have been only two other studies that have explicitly compared phenocopied males to similarly-sized individuals. In the first, conducted by Ewing (1961), the courtship behaviour of males obtained from lines that had been artificially selected for small body size were compared to flies from an unselected population that were reared under crowded conditions. The phenocopied small-bodied males spent more time orientating towards potential mates, but less time vibrating their wings, compared to the control experimentally-evolved small-bodied males (Ewing 1961). Much more recently, Verma *et al*. (2022) compared the effects of males on the fecundity and survivorship of females using flies obtained from populations that has either been selected for fast development and rapid senescence, or from paired ‘control’ lines that had been selected for slow aging from which the ‘selected’ lines had been derived. As these lines had also diverged in body size (control males > selected males), larvae from control lines were grown under crowded conditions to produce smaller phenocopied males closer in size to those of the selected lines (mean male weight (mg) ±SEM control flies: 0.271 ±0.004; selected flies: 0.162 ±0.003; phenocopied-control flies: 0.131 ±0.005, *see Fig S2 in* Verma *et al*. 2022). While they observed that the selected males and the phenocopied males had similar effects on female survivorship and cumulative fecundity, there was considerable heterogeneity in the two small fly types’ effects on female age-specific per capita fecundity – both over time and between experimental blocks – with egg-production in vials containing phenocopied males sometimes considerably greater or considerably lower than in vials that contained control males (*see Fig S3 in* Verma *et al*. 2022). This volatility potentially indicates that the phenocopied flies differed in meaningful and unpredictable ways from the selected males they were being compared to. It is worth noting that in both Ewing (1961) and Verma *et al*. (2022) that phenocopied flies were compared against flies from artificially-selected populations in which body size had evolved, rather than against size-matched flies from their own population, as we did in this study.

While it is beyond the scope of this study to identify the reasons that control and phenocopied males differed in their reproductive successes and effects on females, there are plenty of avenues for speculation and future investigation. Mating success in male flies depends on a number of multiple signal modalities (Hall 1994), whose expression may depend on the specific nature of the developmental environment (*e.g*. Ewing 1961). For instance, differences between phenocopied and control flies may be manifested in the composition of the cuticular hydrocarbons (CHCs) that cover their integument which provide protection against abiotic and biotic environmental stresses, and also play an important role in communication with potential mates. The expression of CHCs in insects can be very plastic (Otte *et al*. 2018), and male fruit flies raised on diets differing in quality can exhibit dramatically different CHC profiles (Bonduriansky *et al*. 2015), thus potentially altering their attractiveness to females. Rearing flies at different densities influences not only the final body size of eclosed flies, but also the size of their internal organs (Mirth and Shingleton 2012), including the primary and secondary organs of the male reproductive system (Morimoto *et al*. 2022). Experimentally increasing larval density leads to progressively smaller males who developed larger testes and ejaculatory ducts, but smaller accessory glands and ejaculatory bulbs (Morimoto *et al*. 2022). Morrow *et al*. (2008) found that the production of sperm cells was affected both by a fly’s genotype as well as the nature of the environment it developed in. Under moderate larval densities flies with genotypes that produced larger body sizes also produced longer sperm, but when the same lines of flies were raised at higher larval densities, the positive correction between body and sperm size disappeared. The disruptive nature of this GxE interaction, means that not only do flies reared at higher densities have smaller sperm, but also that body-size cues about sperm ‘quality’ (*sensu* Pattarini *et al*. 2006) that may be used by females in pre-copulatory mate choice are not reliable. Other components of male ejaculates are also sensitive to developmental conditions. Wigby *et al*. (2016) reported that the effects of experimental manipulation of larval densities were also manifested in differences in the production and transfer of seminal fluid proteins (Sfps), which play an important role in mediating female post-copulatory responses (Sirot *et al*. 2015, Hopkins and Perry 2022). While a positive relationship between male body size and the size of their accessory glands (a site of Sfps production) has been described in groups of flies all reared at the same density (Bangham *et al*. 2002), it is currently unknown how similar the Sfps abundance and/or composition of large/small phenocopied flies are to similarly-sized flies developing under normal conditions. Further differences in the performance and realized success of these males could arise as a result of cryptic female choice exerted by their potential and/or actual mates (Eberhard, 1996), as previous studies have shown that females use the sperm of small and large-bodied phenocopied flies differently (De Nardo *et al*. 2021). The (non-exhaustive) list of possibilities listed above highlight the complex nature of the varied pathways during a fly’s development from genotype to phenotype, and point to the numerous potential difficulties associated with assumptions of using phenocopied flies as substitutes for flies of large or small body size.

If biologists are really interested in understanding the differences in the performance of flies of different body size and/or condition, doing so with individuals who are representative of the population being studied is essential, otherwise we risk making flawed conclusions. It is hoped that the results of this study will lead to the critical reassessment of previous studies that have used phenocopying methods, and going forward, the potential confounds associated with this technique will be clearly acknowledged. In addition to the sieve sorting used in this study, there are numerous new methods of quickly phenotyping flies (*e.g*. Houle *et al*. 2003, Ullah *et al*. 2015) that have developed under evolutionary-relevant conditions, so that one does not need to trade off sample sizes against realism in the pursuit of insight into the operation of sexual selection and conflict.

## Acknowledgements

We would like to thank Yvonne Young, Heather Malek, Garret Bent, Elvira Sathurni, Daniella Balaz, Hanna Dickson, Sarah MacDonald and other the members of the Long Lab for their fly-pushing, behavioural observations and camaraderie. Dr. Scott Ramsay and Natasha Gallo are thanked for their diligent and constructive comments. T.A.F.L. was funded with a Natural Sciences and Engineering Research Council Discovery grant. This work was conducted at Wilfrid Laurier University, which exists on the traditional territory of the Neutral, Anishnawbe, and Haudenosaunee peoples.

